# Influence of Microbiome on Withdrawal-Like Behavior in Planaria

**DOI:** 10.1101/2023.09.12.557338

**Authors:** Enrique Mentado Sosa, John M. Pisciotta, Miguel Guerra-Solano, One Pagan

## Abstract

The microbiome of animals can influence health status, disease susceptibility and can also influence host behavior. This study investigates the effect of microbiome alteration on animal behavior using the planarian model system. For this, *Girardia tigrina* (Brown Planaria) and *Phagocata gracilis* (Black Planaria) were investigated. These flatworms possess a primitive nervous system and exhibit similar addictive and withdrawal behaviors to mammals. Here we tested the hypothesis that alteration of planarian microbiome using the broad-spectrum antibiotics kanamycin and ampicillin could influence the worms’ behavior and withdrawal response to nicotine. After antibiotic treatment period of seven days, the behavior and withdrawal response of four groups of brown and black planaria was analyzed by recording the worms’ motility on a 1 cm^2^ grid. Results support the hypothesis as treatment significantly altered behavior in black worms. The microbiomes of antibiotic treated planaria were analyzed using conventional culture techniques, microscopy and metagenomic methods. Alpha proteobacteria including Sphingomonadaceae were detected in both Brown and Black planarians. This study suggests chemotherapeutic alteration of an animal’s microbiome can influence motility behavior and withdrawal responses to addictive substance and sheds light on the species composition of Planarian microbiomes.

## Introduction

The microbiome of animals plays an increasingly appreciated role in determining the overall health of the host. Mutualistic microorganisms within the gastrointestinal tract of animals aid in the digestion of food and fiber while detoxifying harmful substances (Jing et al. 2020). In so doing they synthesize B group vitamins including niacin and folic acid and other growth factors such as short chain fatty acids and various host essential amino acids (Belzer et al., 2017 Newsome et al., 2020). The microbiome of animals, including humans, influences host immunity and can affect the central nervous system which regulates behavior (Yadev et al., 2017). For instance, the microbiome is needed to stabilize circadian rhythm at the transcriptome level in Drosophila model organism system (Zhang et al. 2023).

Planarian flatworms are widely utilized as model organisms to study animal behavior, pharmacological responses and tissue regeneration (Ivankovic et al., 2019, Mohammed et al., 2018). *Girardia tigrina* possess powerful regenerative abilities, which enables the worm to regenerate body parts and organs from existing pluripotent stem cell promoters within tissues (de Oliveira et al. 2018). Planarian flatworms demonstrate a complex ability to learn and problem-solve and can completely regenerate their brain and central nervous system if damaged (Morokuma, et al 2017). Administration of Bacillus-based bioinsecticides have recently been shown to alter behavior and regeneration rates planaria (Silva, et al. 2022). This suggests the microbiome could play an important role in the modulation of behavior in response to chemical, such as pharmacological agents.

Here we hypothesize that the alteration of planarian worms’ microbiomes using broad-spectrum antibiotics will alter the behavior of the Planarians and their response to withdrawal from the addictive substance nicotine. We further hypothesize that differences in microbiomes exist between different planarian genera. This research can help shed light on how the microbiome may influence behavior in animals while enhancing our understanding of the microbial ecology of disparate planarian flatworms.

## Materials and Methods

### Antibiotic Treatment and Behavioral Analysis

Eight sets of worms were prepared for behavioral analysis. These consisted of Brown worms with no drug, Brown worms with Ampicillin, Brown worms with Kanamycin, Brown worms with Ampicillin + Kanamycin. The same four treatment groups were similarly carried out with Black worms. After seven days of drug treatment, worms were exposed to four different conditions before conducting motility testing on a 1cm^2^ grid for five minutes. The four room temperature conditions were: One hour in nicotine, then motility testing in nicotine, one hour in nicotine, then motility testing in artificial pond water, one hour in artificial pond water, then motility testing in nicotine. The no nicotine control group was one hour in artificial pond water, then motility testing in artificial pond water.

### Culture Based and Microscopic Analysis

Following the seven-day antibiotic treatment period, worms from each group were weighed then homogenized in artificial pond water using sterile, disposable mortar and pestles. The homogenates were serially diluted then spread plated onto sterile Brain Heart Infusion (BHI) agar plates and incubated at room temperature for seven days. After quantifying the number of colonies on each plate, Gram stained slides of four representative microbes from Brown and Black worm BHI plates were generated and visualized using microscopy.

### Metagenomic DGGE Analysis

DNA was extracted from each of the eight worm homogenates using the PowerBioFilm® DNA Isolation Kit (Mo Bio Inc.). The isolated DNA from each worm group was amplified using Polymerase Chain Reaction (PCR) and 16s primers (338f + 907rev). Following amplification, a 70-40% DGGE gel was run overnight at 90V then stained with Sybr-gold for analysis of differences in microbial communities in the different treatment groups.

### Preparation of DNA Library

DNA was extracted from each of the worm homogenates was subjected to a quality check and quantification with a NanoDrop spectrophotometer. Each of the eight samples of extracted DNA was adjusted to 10 ng of Genomic DNA with PCR-grade water was added to adjust the volume appropriately. The 16s Barcodes for each sample 1-8 were thawed at room temperature, then each tube was individually mixed. The eight barcoded samples were amplified using the conditions specified in the 16s Barcoding Kit protocol. PCR amplified samples were subsequently transferred into fresh, sterile tubes and 30ul of AMPure XP magnetic beads were added and incubated on a rotator mixer for 5 minutes at room temperature. Tubes were spun down at 10,000 g for 30 seconds, then placed on a magnet to immobilize beads as supernatant was removed. Next, 200ul of 70% ethanol was added to each tube then removed to wash. Barcoded samples were spun down at max speed for 30 seconds then placed onto a magnet. Residual ethanol was removed and the pellets were air dried prior resuspension in 10ul of 10mM Tris-HCl pH 8.0 with 50mM NaCl at room temperature for 2 minutes. Ten (10) ul of clear eluate from each of the eight tubes was retained in eight, separate 1.5ml Eppendorf tubes. Into a single 1.5ml Eppendorf tube, each of the eight barcoded libraries were pooled in the appropriate ratios to a total of 100 femptomoles, and 10mM Tris-HCl pH 8.0 with 50mM NaCl was added reach a total volume of 10ul. 1ul of RAP was added the pooled library, and the tube was mixed gently and incubated at room temperature for 5 minutes.

### MinION 16s rDNA Sequence Analysis

The Sequencing Buffer (SQB), Loading Beads (LB), Flush Tether (FLT) and one tube of Flush Buffer (FB) were thawed at room temperature then kept on ice. The Sequencing Buffer (SQB) and Flush Buffer (FLB) tubes were mixed individually by vortexing and spun down. Once mixed, each tube was kept on ice. The MinION nanopore DNA sequencer was opened, and the flow cell was placed inside and primed and loaded as per the manufacturer’s instructions. The Sequencing Buffer (SQB) and Loading Beads (LB) were mixed individually by pipetting and then 70 ul of the pooled library was loaded and the samples sequenced and recorded using a 1 TB SSD hard drive computer. Once complete, metagenomic sequence data was analyzed via the EPI2ME online database.

## Results

Untreated blank and brown planarians did not differ with regard to their motility rates (Fig 1). Motility of black planarian worms was significantly reduced by antibiotic treatment compared to no drug control (Table 1). Conversely, brown worm behavior was not significantly affected by antibiotic treatment. Since black worms were significantly affected by antibiotic treatment, further behavioral tests involving nicotine withdrawal were conducted using brown worms. These results suggest microbiome modification alters withdrawal-like behavior considering that the untreated worms showed elevated motility (Table 1).

**Figure 1.**
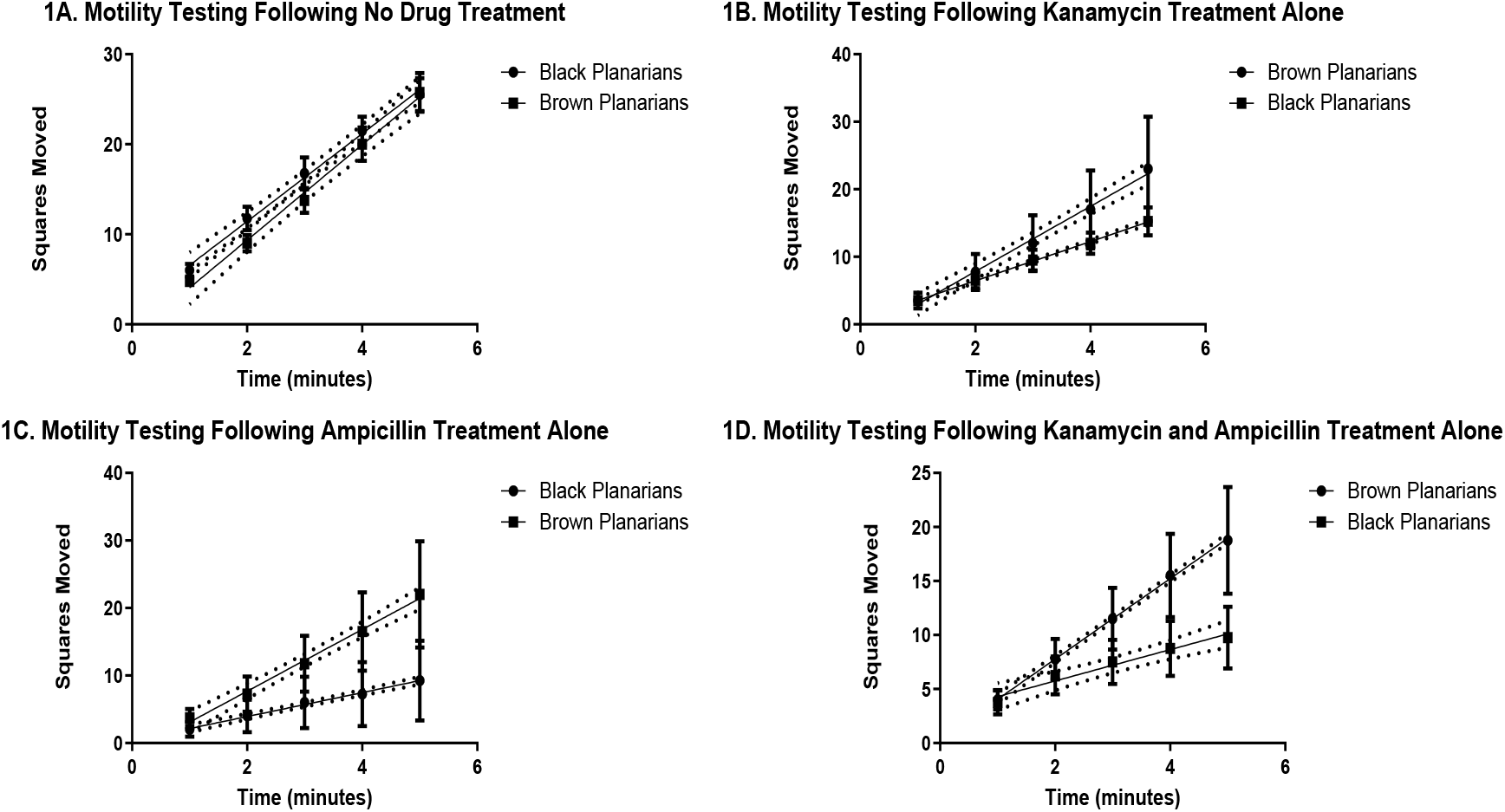
Brown and Black Worm Motility Test Following Antibiotic Treatments.

**Table 1.**
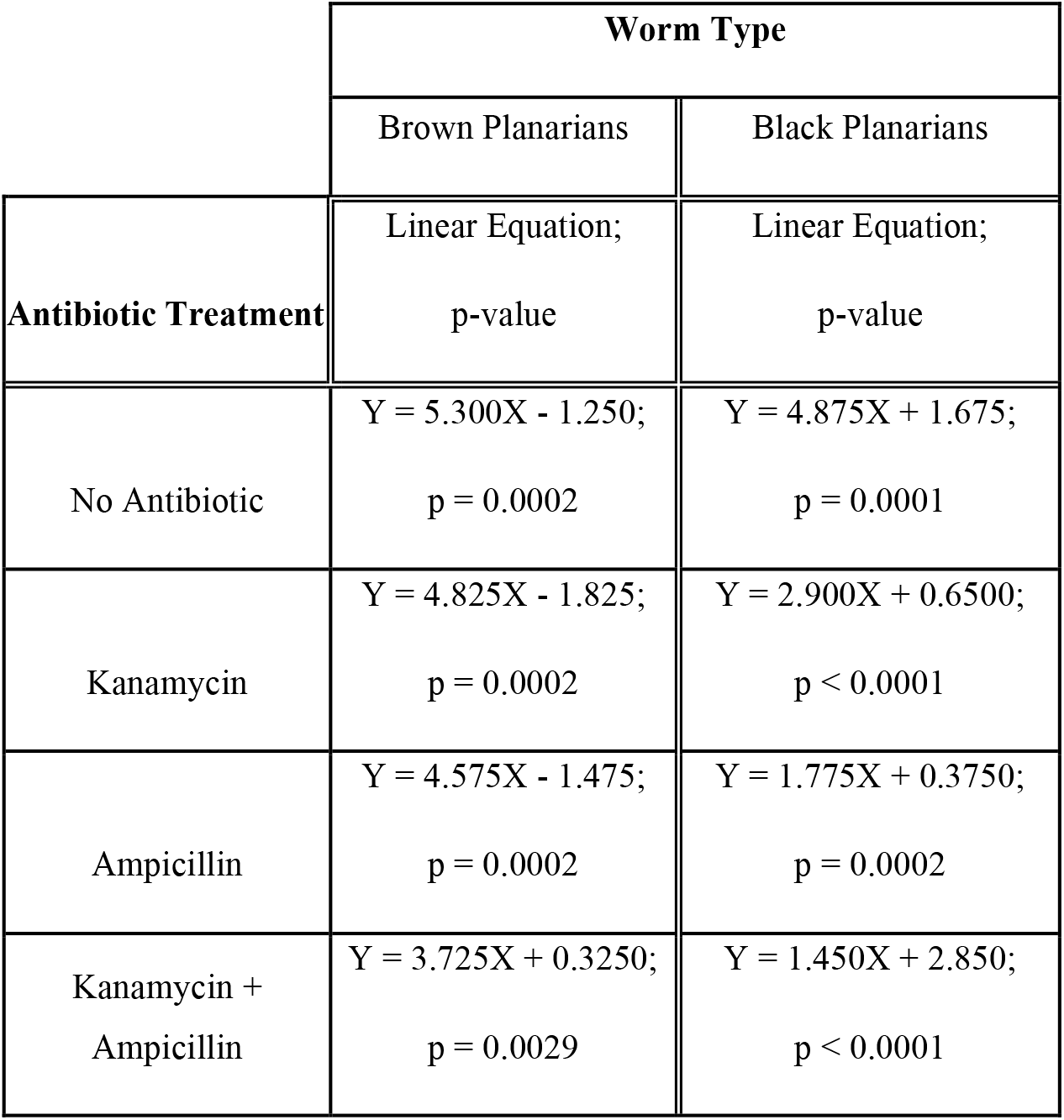
Brown and Black Worm Motility Test Statistics.

|  | Worm Type |  |
| --- | --- | --- |
|  | Brown Planarians | Black Planarians |
| Antibiotic Treatment | Linear Equation;<br>p-value | Linear Equation;<br>p-value |
| No Antibiotic | $Y = 5.300X - 1.250$;<br>$p = 0.0002$ | $Y = 4.875X + 1.675$;<br>$p = 0.0001$ |
| Kanamycin | $Y = 4.825X - 1.825$;<br>$p = 0.0002$ | $Y = 2.900X + 0.6500$;<br>$p < 0.0001$ |
| Ampicillin | $Y = 4.575X - 1.475$;<br>$p = 0.0002$ | $Y = 1.775X + 0.3750$;<br>$p = 0.0002$ |
| Kanamycin +<br>Ampicillin | $Y = 3.725X + 0.3250$;<br>$p = 0.0029$ | $Y = 1.450X + 2.850$;<br>$p < 0.0001$ |

**Figure 2.**
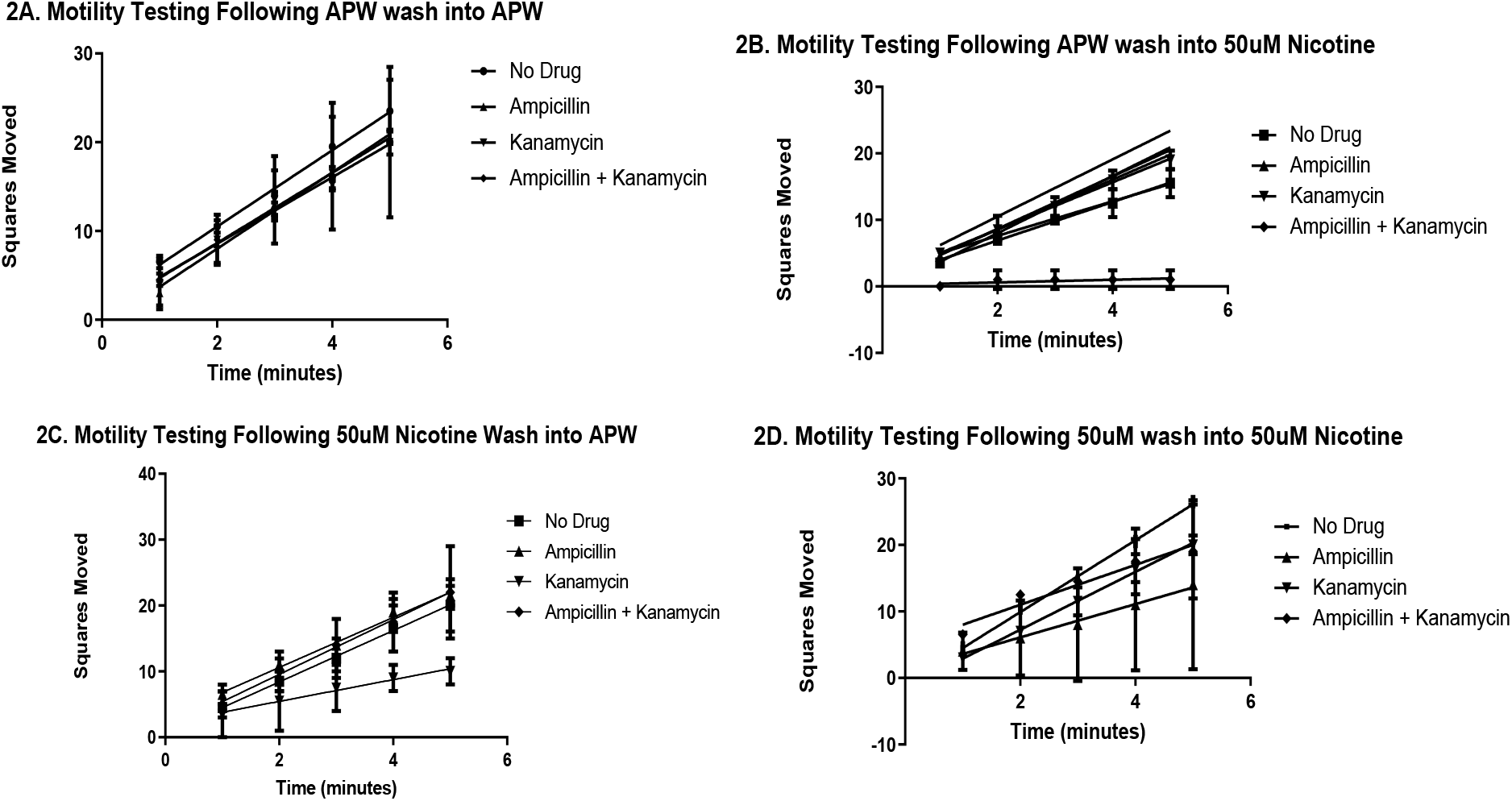
Brown Worm Behavioral Motility Test Following Antibiotic Treatments.

Bacterial colonies were obtained from diluted, spread plated Brown and Black worm homogenates grown on Brain Heart Infusion (BHI) Agar for 7 days at room temperature. These isolates demonstrated morphologically distinct bacterial colonies between the two worm species (Fig 3, Fig 4). Brown worms demonstrated a larger number of Kanamycin resistant bacteria (54 vs 32 colonies). Black worms demonstrated a larger number of Ampicillin resistant bacteria (51 vs 8 colonies). Optical microscopy of Gram stains indicated gram negative rods (Figs 3D and 4D).

**Figure 3.**
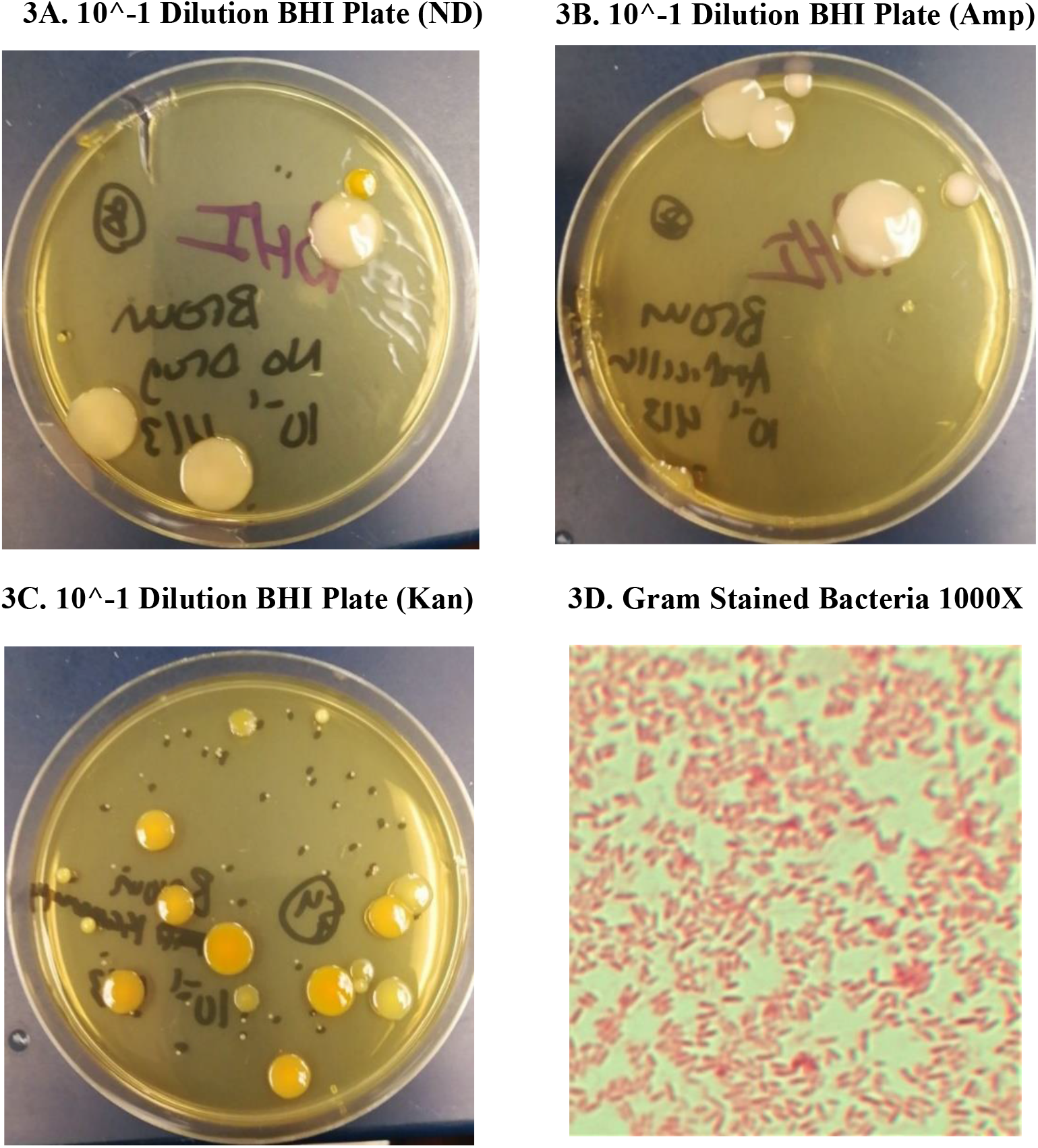
Brown Worm Bacteria on BHI Agar.

**Figure 4.**
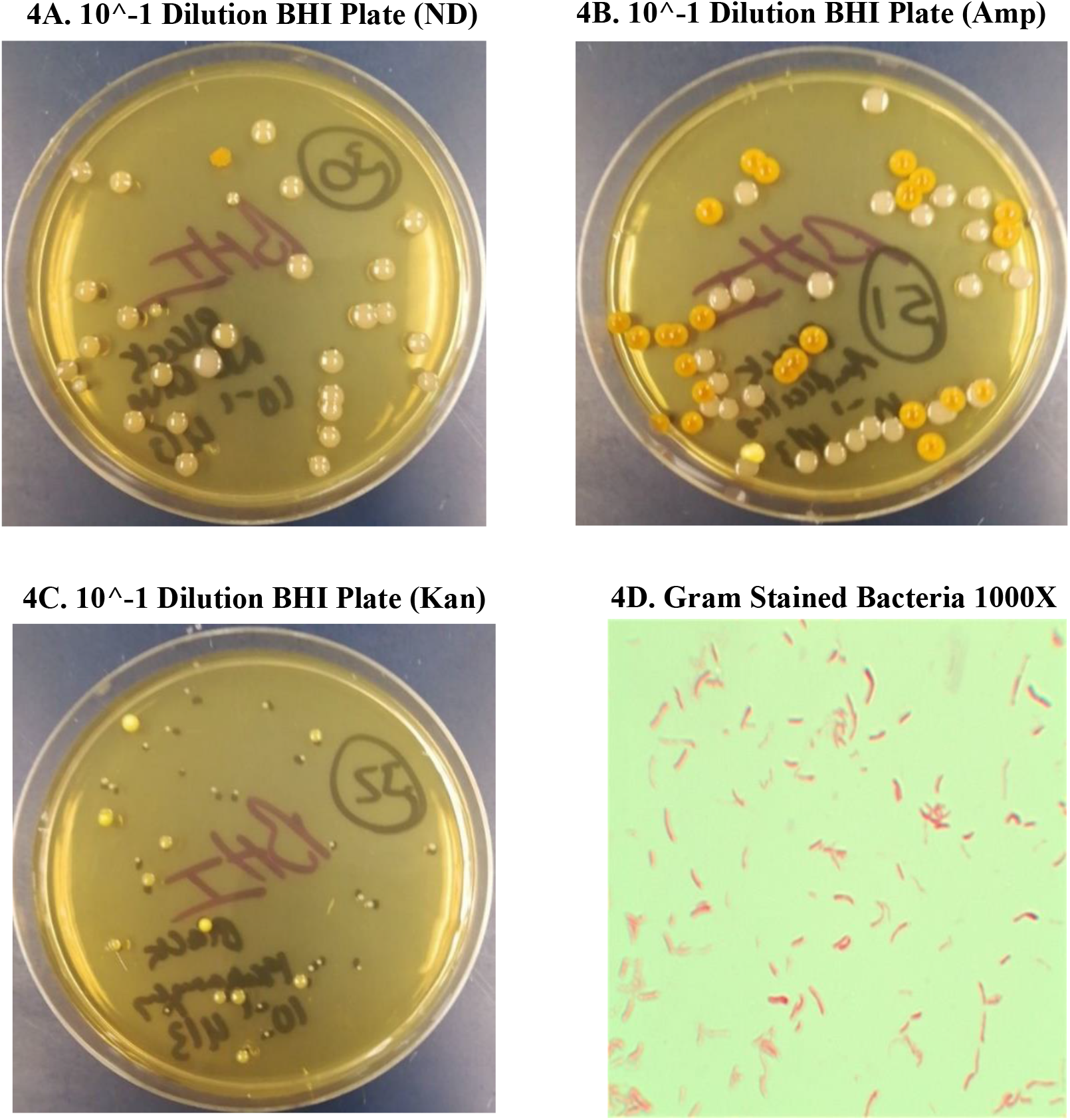
Black Worm Bacteria on BHI Agar.

Metagenomic 16s rRNA gene DGGE analysis of antibiotic treated worms demonstrated that antibiotic treatment reduced the overall diversity of bacteria in both worm types (Fig. 5). The red box demonstrates a larger diversity of GC rich bacteria in non-treated Brown worms compared to drug treated Brown worms. The green squares demonstrate a similar response outcome to drug treatment between non-treated Black worms and drug-treated Black worms.

**Figure 5.**
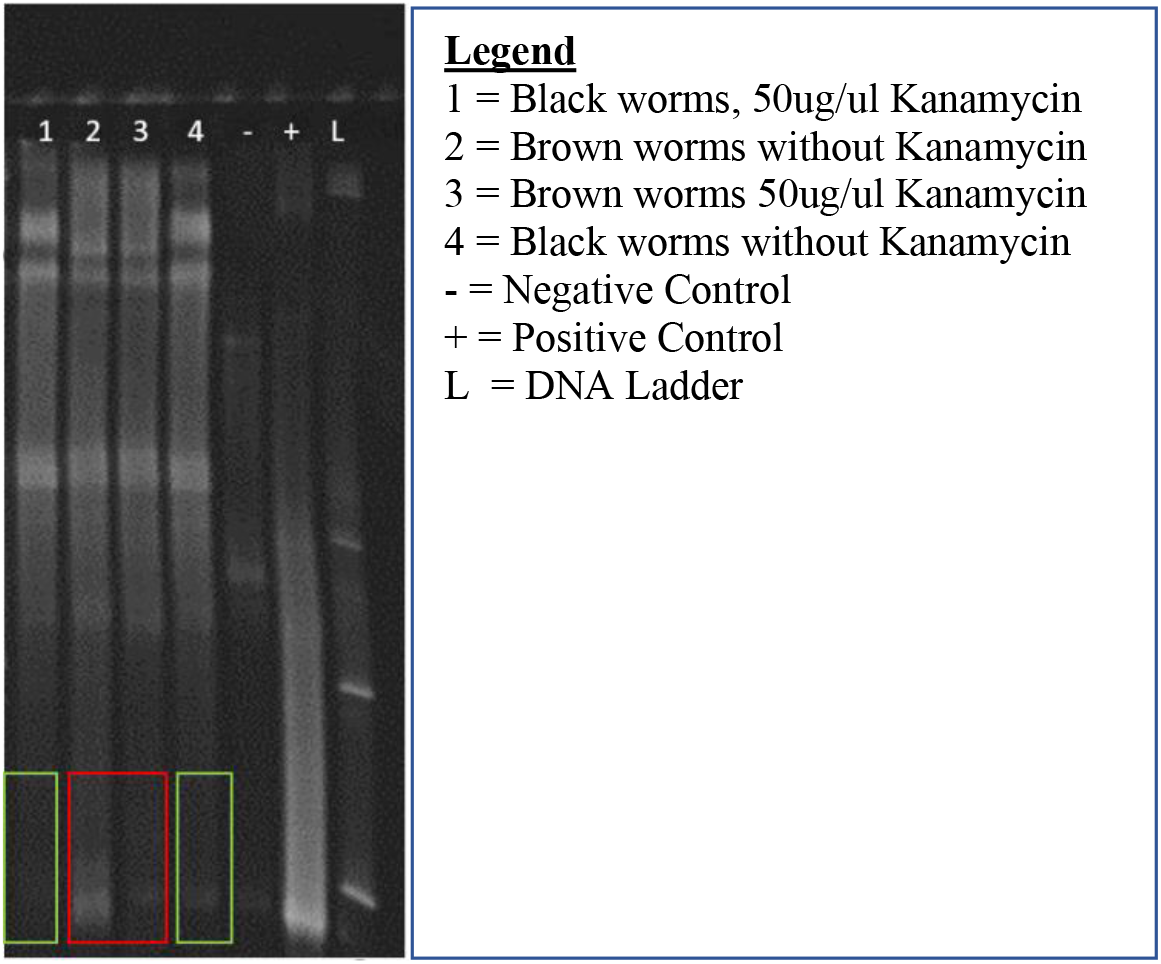
Metagenomic DGGE Community Analysis of Brown and Black Worms Following Treatment with Kanamycin.

Metagenomic DNA 16s rDNA gene sequence analysis of Brown and Black worm barcoded libraries yielded 189 total reads. The most common bacteria in each worm type were for Non-treated Brown worms Edaphobacter, Dyella, and Cutibacterium (Fig 6). For ampicillin treated Brown worms it was Sphingomonadaceae family members. Kanamycin treated Brown worms contained Streptococcus and Pedobacter. Ampicillin and Kanamycin Brown worms contained Acidobacteriaceae, Rhodanobacteraceae, and Oxalobacteraceae. Thus, the dual treatment did not fully ablate the bacterial microbiome.

**Figure 6.**
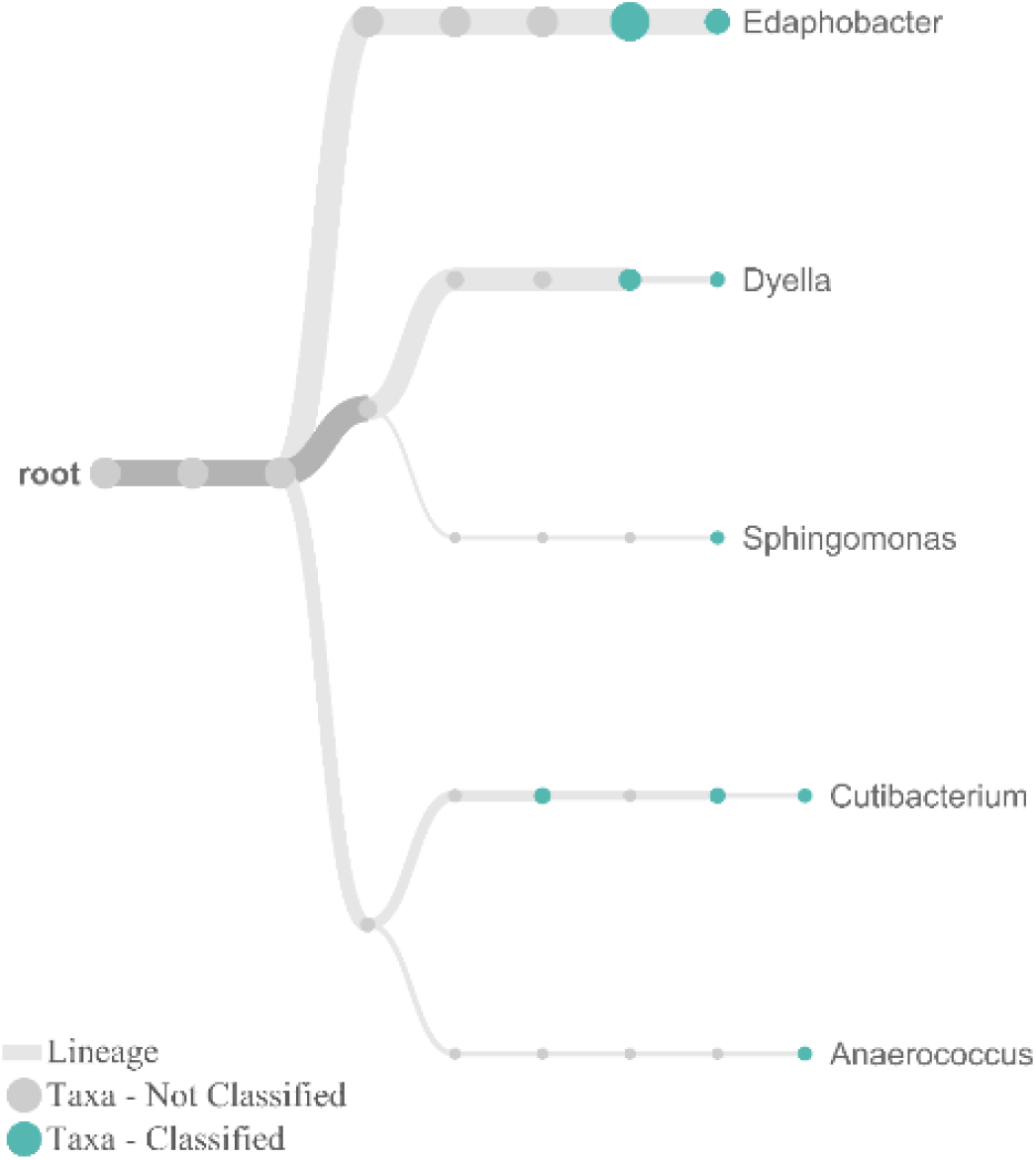
Taxonomic Tree of Non-treated Brown worm Microbiome.

Non-treated Black worms contained detectable Sphingomonas, and Pedobacter, similar to Brown worms, suggesting the microbiome of different genera of planarians overlaps. No reads were detected by MinION for the ampicillin treated Black worms although kanamycin treated Black worms indicated Pedobacter, Sphingomonas, and Mucilaginbacter. No amplicon reads were detected by the MinION for the ampicillin and kanamycin treated Black worms. This further support the notion that antibiotic based ablation of the host microbiome can deleteriously affect host motility (Fig 1D).

## Conclusions

Based on the results of this study, it appears that the black worm, *P. gracilis*’ motility and microbiome composition are adversely affected by antibiotic-mediated perturbation of the microbiome to a greater degree than brown worms (Fig 1, Table 1). Untreated worms demonstrated highest motility, suggesting an intact microbiome may influence motility and possibly withdrawal-like behavior. Silva found that addition of certain bacteria, namely the gram positive rod *Bacillus thuringiensis kurstaki*, significantly diminished *G. tigrina* motility (Silva et al. 2022). It is therefore possible that removal of certain microbiome populations via administration of broad spectrum antibiotics could similarly alter motility. Certain commensal bacteria have been found to produce metabolites that affect the host worm in deleterious ways, at the transcriptional level (William et al 2020). It is plausible that selective disruption of some microbiome species could open niche space for drug-resistant, detrimental species. Indeed, in human’s treated with broad spectrum antibiotics opportunistic bacterial pathogens such as *Clostridium difficile*.

Interestingly, antibiotic treatment did not have a significant effect on withdrawal-like behavior in the brown worms (Fig 2., Table 2). This might be due to populations of innately antibiotic resistant bacteria associated with brown worms. Culture based analysis (Table 3-4, Fig 3-4) and DGGE prfiling suggest differences in ampicillin and kanamycin resistant bacteria ma exist between worm species.

**Table 2.** Brown Planaria Motility Test Statistics.

|  | **Behavior Treatment Conditions** |  |  |  |
| --- | --- | --- | --- | --- |
|  | APW Wash into APW | 50uM Nicotine Wash into APW | APW Wash into 50uM Nicotine | 50uM Wash into 50uM Nicotine |
| **Antibiotic Treatment** | Linear Equation; p-value | Linear Equation; p-value | Linear Equation; p-value | Linear Equation; p-value |
| No Antibiotic | $Y = 4.300X + 1.900$; $p < 0.0001$ | $Y = 3.900X + 0.6000$; $p = 0.0006$ | $Y = 2.900X + 1.100$; $p < 0.0001$ | $Y = 5.400X - 0.9000$; $p < 0.0001$ |
| Kanamycin | $Y = 3.950X + 0.7500$; $p = 0.0040$ | $Y = 1.650X + 2.150$; $p = 0.0777$ | $Y = 3.550X + 1.450$; $p < 0.0001$ | $Y = 4.350X - 1.450$; $p < 0.0001$ |
| Ampicillin | $Y = 4.300X - 0.600$; $p < 0.0001$ | $Y = 3.800X + 3.000$; $p < 0.0001$ | $Y = 2.600X + 2.400$; $p < 0.0001$ | $Y = 2.500X + 1.100$; $p = 0.1404$ |
| Kanamycin + Ampicillin | $Y = 3.750X + 1.050$; $p < 0.0001$ | $Y = 4.150X + 1.250$; $p = 0.0058$ | $Y = 0.2000X + 0.2000$; $p = 4.186$ | $Y = 3.000X + 5.000$; $p = 0.0037$ |

Metagenomic DGGE analysis of brown and black worms indicated that differences in microbial communities exist in different planarian genera. Furthermore, antibiotic treatment alters gut bacteria population in worms (Fig 5). 16s rDNA gene sequence analysis confirmed bacterial populations within brown and black treatment groups. In this study, Pedobacter was observed in both planarian genera. Pedobacter, it appears, is a broadly conserved constituents of the microbiome in disparate Planaria flatworms as Pedobacter was also detected in the European planarian *Schmidtea mediterranea* (Kangale, 2020, Kangale 2021). Collectively these results indicate antibiotic treatment affects the planarian microbiome and that this disruption may alter behavior and behavioral responses to addictive substances. Future studies will aim to investigate the mechanistic basic through which the animal microbiome may influence animal host behaviors.

## Acknowledgements

The WCU Biology Dept. and College of Science and Mathematics provided funding for this research study.

