## Supplementary figures and images for "Influence of Microbiome on Withdrawal-Like Behavior in Planaria"

### supplemental data

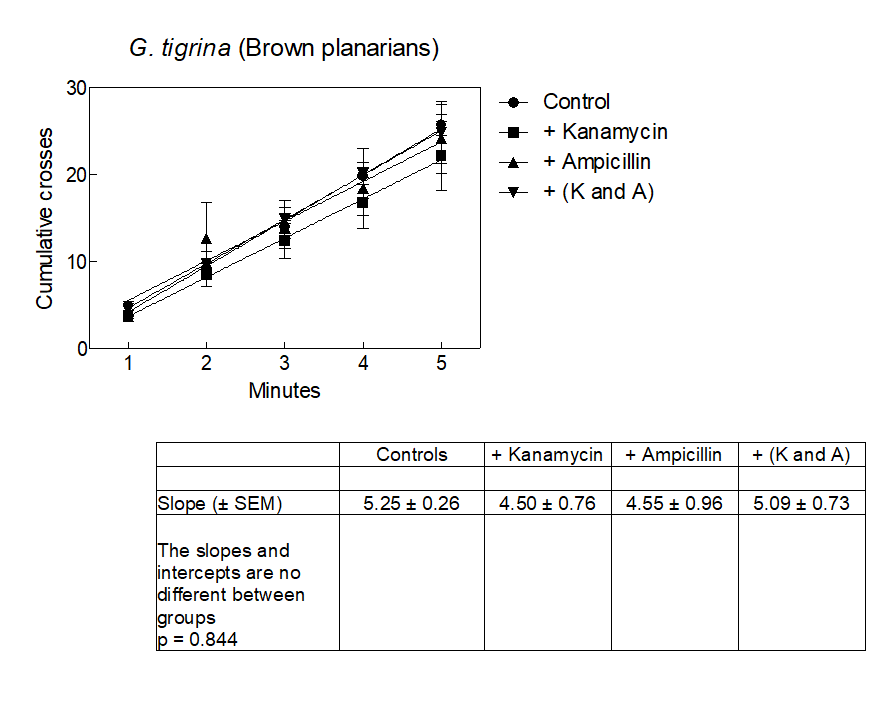

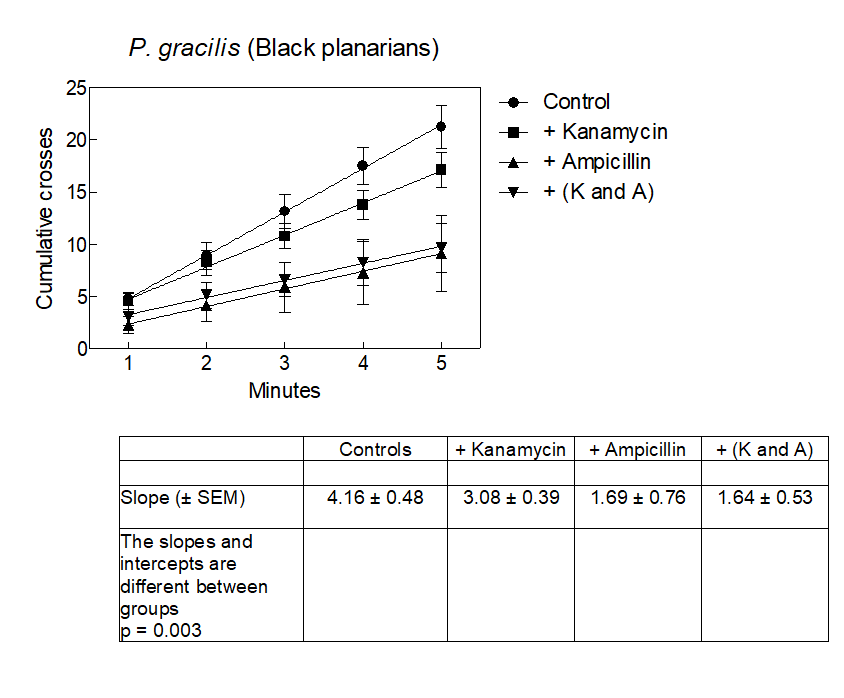
